# Truncating Variant Burden in High Functioning Autism and Pleiotropic Effects of *LRP1* Across Psychiatric Phenotypes

**DOI:** 10.1101/429779

**Authors:** Bàrbara Torrico, Alex D Shaw, Roberto Mosca, Norma Vivó-Luque, Amaia Hervás, Noèlia Fernàndez-Castillo, Patrick Aloy, Monica Bayés, Janice M. Fullerton, Bru Cormand, Claudio Toma

**Author notes:** These authors contributed equally (Bru Cormand, Claudio Toma). **Corresponding authors:** Claudio Toma, Neuroscience Research Australia, Margarete Ainsworth Building, Barker Street, Randwick NSW 2031, Sydney, Australia., Bru Cormand, Departament de Genètica, Facultat de Biologia, Universitat de Barcelona, Prevosti Building, floor 3, Av. Diagonal 643, 08028 Barcelona, Spain.

## Abstract

Previous research has implicated *de novo* (DN) and inherited truncating mutations in autism spectrum disorder (ASD). We aim to investigate whether the load of inherited truncating mutations contribute similarly to high functioning autism (HFA), and to characterise genes harbouring DN variants in HFA.

We performed whole-exome sequencing (WES) in 20 HFA families (average IQ = 100). No difference was observed in the number of transmitted versus non-transmitted truncating alleles to HFA (117 vs 130, *P* = 0.32). Transmitted truncating and DN variants in HFA were not enriched in GO or KEGG categories, nor autism-related gene sets. However, in a HFA patient we identified a DN variant in a canonical splice site of *LRP1*, a post-synaptic density gene that is a target for the FMRP. This DN leads to in-frame skipping of exon-29, removing 2 of 6 blades of the β-propeller domain-4 of LRP1, with putative functional consequences. Results using large datasets implicate *LRP1* across psychiatric diseases: i) DN are associated with ASD (*P* = 0.039) and schizophrenia (*P* = 0.008) from combined sequencing projects; ii) Common variants using Psychiatric Genomics Consortium GWAS datasets show gene-based association in schizophrenia (*P* = 6.6E-07) and across six psychiatric diseases (meta-analysis *P* = 8.1E-05); and iii) burden of ultra-rare pathogenic variants is higher in ASD (*P* = 1.2E-05), using WES from 6,135 schizophrenia patients, 1,778 ASD patients and 6,245 controls. Previous and current studies suggest an impact of truncating mutations restricted to severe ASD phenotypes associated with intellectual disability. We provide evidence for pleiotropic effects of common and rare variants in the *LRP1* gene across psychiatric phenotypes.

## Introduction

Autism spectrum disorder (ASD) is characterized by impairments in social interactions, communication and repetitive behaviours. A prevalence of 1% in general population makes ASD one of the most prevalent disorders in childhood (1). The clinical phenotype is heterogeneous, and includes a broad range of comorbidities such as epilepsy, language impairment, anxiety, sleep disorders or attention-deficit hyperactivity disorder (ADHD) (2). However, one of the most remarkable clinical features in ASD is represented by intellectual disability (ID), which is present in a considerable proportion of patients (3), and associated with the most severe phenotypic outcomes across the spectrum (4). ASD patients with higher intelligence quotient (IQ>70) and average or high cognitive abilities are often considered in a more homogenous clinical group referred to as high functioning autism (HFA).

Recent family studies confirm that genetic factors play a considerable role in ASD. Heritability (*h^2^* = 0.8) is one of the highest amongst neuropsychiatric disorders, with environmental factors minimally involved in disease risk (5). Despite the high heritability, the specific genetic risk factors remain largely unknown, and only a small proportion of the approximately 1,000 genes estimated to be involved in ASD have been identified (6, 7).

The biological processes involved in ASD are yet to be fully determined, although evidence suggests that ASD may have a neuroinflammatory base (8), result from mitochondrial energetic deficits (9, 10), or considered a synaptopathology (11).

Genetic studies conducted over the last two decades converge on a genetic model in which both multiple common variants of small effect size and a discrete number of rare variants with higher penetrance, shape ASD genetic liability. The first genome-wide association studies (GWAS) pointed at several common variants in ASD (12–14), which were not replicated in a recent well-powered study of European populations (15). Although common risk variants are estimated to explain approximately 20–50% of the genetics in ASD (16, 17), larger cohorts are needed to identify individual allelic contributions (15). Also the last efforts combining several large international GWAS datasets in a meta-analysis failed to identify genome-wide significant hits (18).

Larger sample sizes and increased sensitivity in whole-exome or whole-genome sequencing studies (WES or WGS) have exponentially increased the identification of novel risk genes, setting the time to resolve most of the missing heritability in psychiatric diseases. Several sequencing studies have implicated both rare *de novo* variants (DNs) and rare inherited single nucleotide variation (SNVs) in ASD (19–21). The first WES studies implicated DN point mutations in disease pathogenesis in singleton ASD families (22–25), estimating that this mutation class may explain between 5–20% of genetic liability (23, 26, 27). DN variants are considered highly deleterious, and an excess of DN truncating gene variants was found in autistic probands (24). Interestingly, several independent studies found a correlation between the higher burden of DN truncating variants and lower IQ in ASD (22, 23, 25). Our group was the first to assess the impact of inherited rare variants in multiplex ASD families (28), which suggested that apart from DN, inherited truncating variants also play a significant role in disease pathogenesis with a higher burden in ASD patients (28). The contribution of inherited truncating variants in ASD was replicated later in a large sample by comparing probands and their unaffected siblings (29).

However, it is still unclear whether inherited truncating mutation burden is etiologically relevant in all autistic patients or if it is prominent only in those severe ASD cases associated with intellectual disability. Therefore, in this study we aimed to: i) determine whether rare inherited truncating mutations are distinctly implicated in high functioning autism; ii) identify molecular pathways or biological categories in ASD by considering the entire pool of severe mutations, including DN and inherited truncating variants; and iii) identify potential novel candidate genes for ASD.

## Results

### Inherited truncating alleles in high functioning autism

A total of 247 truncating alleles were validated after whole exome sequencing (WES) in 20 HFA families (47% indel-frameshift, 37% nonsense, 16% canonical splice-site variants) and are listed in supplementary material, Table S1. We assessed whether the number of truncating alleles transmitted to HFA probands was higher than those non-transmitted, and found no significant difference across all families (z = 0.992, *P* = 0.32) (Table 1), nor when considering each individual family (*P*>0.05). Moreover, no difference was observed after restricting analysis to truncating alleles in brain-expressed genes (z = 0.207, *P* = 0.84) (Table 1). A similar ratio of transmitted/non-transmitted alleles was observed in unaffected siblings (Table 1).

**Table 1.**
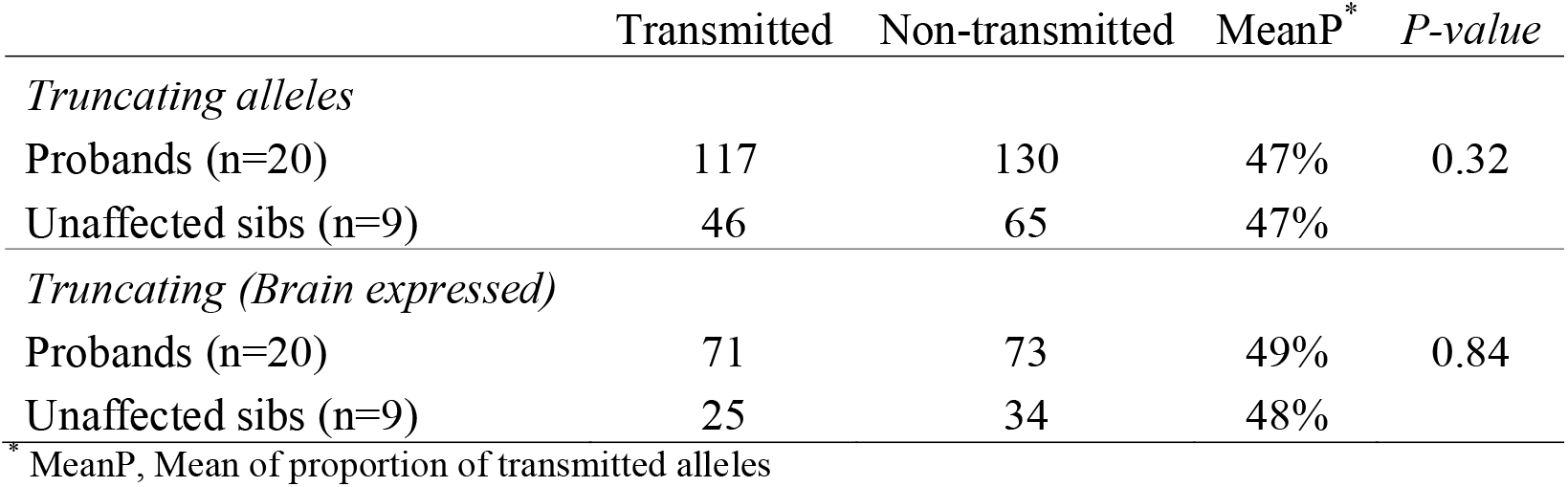
Transmitted and non-transmitted truncating alleles found in 20 singletons families with HFA.

### Identification of de novo variants in HFA

We examined the impact of *de novo* (DN) variants across the 20 HFA families and 16 variants were validated: 13 missense, 2 synonymous and 1 splicing variant (supplementary material, Table S2). Missense variants, predicted pathogenic by both SIFT and PolyPhen-2, were found in *PCDH15, ADD3, GALNT6* and *TEX14*, and a potential functional DN variant was found in a canonical acceptor splice site in *LRP1* gene in one proband (Supplementary material, Fig. S1).

### Enrichment analysis of inherited truncating and DN variants

We performed an enrichment analysis for Gene ontology (GO) categories and KEGG pathways of the combined pool of highly damaging variants including both inherited truncating alleles and DN variants found in the 20 HFA probands. Although categories with a Torrico et al plausible role in ASD were found in the top hits (Brain morphogenesis, *P* = 0.001; neurotransmitter receptor complex, *P* = 0.003; histidine metabolism, *P* = 0.023), none were significant after multiple correction (supplementary material, Table S3 and S4).

Enrichment analysis against gene-sets previously implicated in psychiatric disorders (30, 31) found no evidence of enrichment (Table 2). However, the *LRP1* gene was present in all gene-sets under study, namely: the post-synaptic density (PSD) genes, DN variants in ASD, DN variants in schizophrenia and FMRP target genes (Table 2), warranting further investigation of this candidate gene.

**Table 2.** Gene-set enrichment analysis of transmitted truncating alleles and *de novo* variants in the 20 HFA probands (130 genes). The same analysis was performed in the non-transmitted truncating alleles (128 genes) (P>0.05).

| Gene set<br>(n genes) | O (E) | Empirical *P*-value | Genes |
| --- | --- | --- | --- |
| PSD<br>(1435) | 6 (9.9) | 0.95 | - *TBC1D24*, *UGP2*, *ADD3*, *CDH2*,<br>*GRIN2B*, ***LRP1*** |
| FMRP targets<br>(835) | 4 (6.4) | 0.91 | - *GRIN2B*, *NPAS2*, *DIDO1*, ***LRP1*** |
| *De novo* AUT<br>(768) | 8 (7.2) | 0.43 | - *METTL16*, *GRIN2B*, *EFCAB5*,<br>*CACNA1S*, *MUC17*, *UTP20*, ***LRP1***,<br>*DNAH11* |
| *De novo* SCZ<br>(694) | 9 (7.1) | 0.25 | - *ZNF551*, *FILIP1*, *IGSF22*, *CLTCL1*,<br>*MYH7B*, *CACNA1S*, *MUC17*,<br>*VPS13C*, ***LRP1*** |

Abbreviations: DN, *de novo* variants; O: number of genes observed in this category; E, number of genes expected in this category; PSD, genes expressed in the Post-Synaptic Density (57); FMRP, Fragile X mental retardation protein target genes (58); *De novo* AUT, de novo variants found in autism (30); *De novo* SCZ, de novo variants found in schizophrenia (59); The recurrent gene *LRP1* is in bold.

### LRP1 de novo splice site variant leads to skipping of exon 29 with functional consequences

The *LRP1* DN variant found in proband SJD_33.3 (chr12:57573110, A/G) is located in a highly conserved canonical splice site and it is absent from the *gnomAD* database (http://gnomad.broadinstitute.org/). The AG to GG change is predicted to be functional and highly deleterious according to three splicing-based analysis tools: MaxEntScan (score = 4.12), SPANR (dPSI-wt: 0; dPSI-mut: –12.95), and finally HSF (wt score: 86; mut score: 57), predicting a disrupted acceptor site of exon 29 (Figure 1A-B). Examination of *LRP1* mRNA from the patient’s blood lymphocytes confirmed an in-frame skipping of exon 29 (Figure 1C). The expression levels of transcripts from the mutated and WT alleles were similar (details in the Supplement). The *LRP1* DN variant was also present in the patient’s buccal cells from a saliva sample, which indicates a germline mutational origin and the presence of the abnormal transcript in all tissues where *LRP1* is expressed, including brain. Moreover, this DN mutation was absent in two additional unaffected siblings from this family.

To assess the potential functional consequences of exon 29 skipping (p.1580–1655del) at protein level, we performed a modelling study of the LRP1 protein domains (NP_002323.2). Exon 29 encodes part of an YWTD β-propeller domain of LRP1 (Figure 2A). Each of the β-propeller domains is the result of six YWTD repeats that are organized in six four-stranded β-sheets (blades) arranged radially about a central symmetry axis. We generated a structural model of the third and fourth LRP1 β-propeller domains, based on the homology of LRP6, which showed that exon 29 encodes the first two blades of the fourth β-propeller domain (Figure 2B). When we modelled the mutated form of LRP1, lacking the exon 29, the sequence alignment matched the β-propeller domains 3 and 5, whereas the β-propeller domain 4 segment was unaligned and cannot be folded into a globular structure. However, the poor quality of the model does not exclude the possibility that the mutated β-propeller domain 4 may fold into an ordered structure.

**Figure 1:**
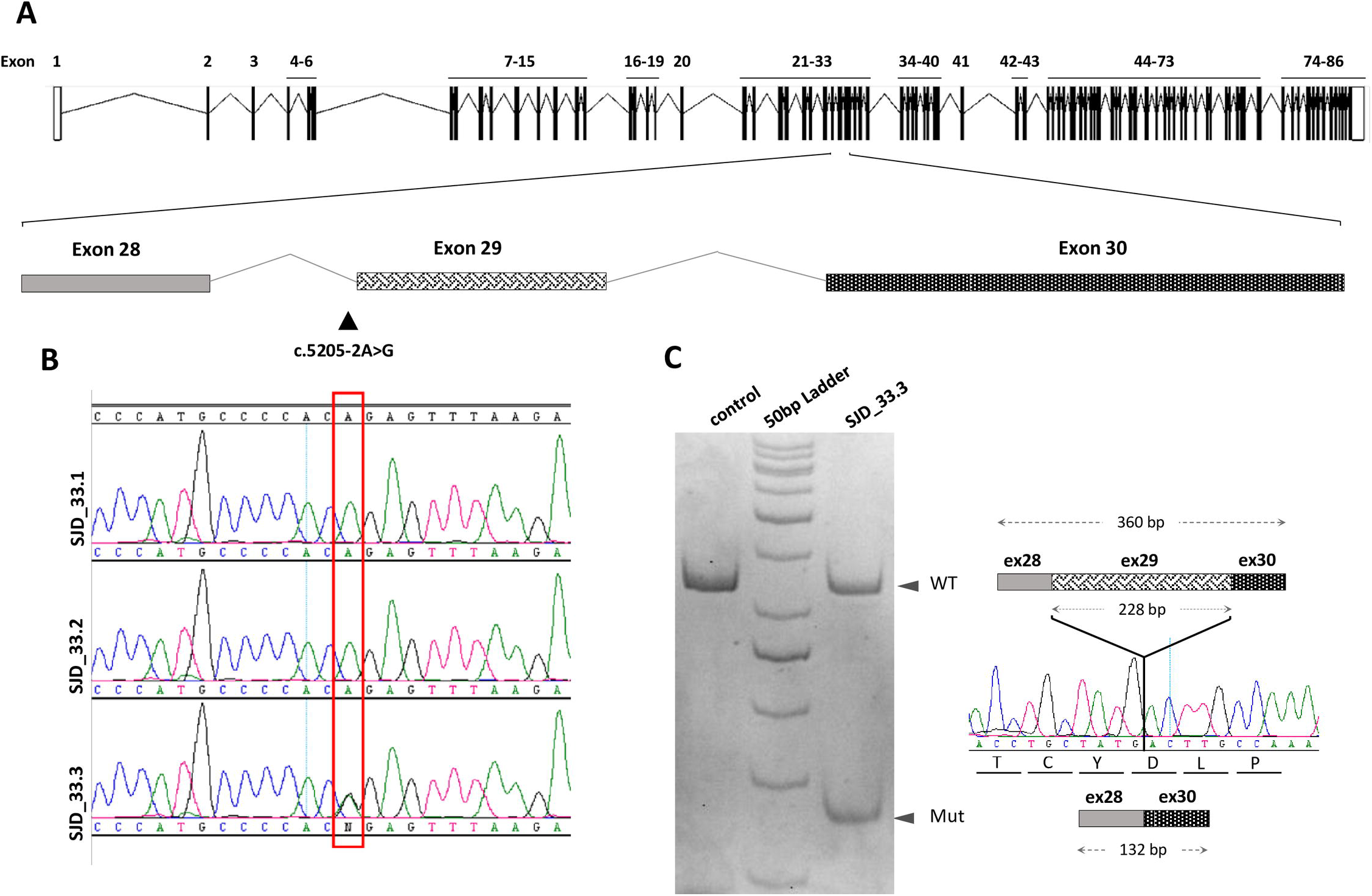
Effect of the identified DN change on *LRP1* splicing. **A)** Schematic structure of *LRP1* gene (NM_002332), with exons from 28 to 30 amplified below. The mutation site is indicated by a triangle (c.5205–2A>G). **B)** Sanger validation of the variant identified by whole exome sequencing in the SJD_33.3 proband, which position is framed by a red box. **C)** PCR analysis of *LRP1* cDNA from lymphocytes visualized on polyacrylamide gel. For the wild type (WT) transcript, the PCR amplicon of 360bp included a fragment of exon 28, the entire exon 29 and a fragment of exon 30, whereas the mutated transcript (Mut) generated a smaller fragment (132bp) lacking exon 29 (76 amino acids), which generate an in-frame transcript showed in figure. The two transcripts spanning exon 28 to 30 are represented with the sequenced smaller band of the mutated allele. SJD_33.1: proband’s father; SJD_33.2: proband’s mother; SJD_33.3: proband.

**Figure 2:**
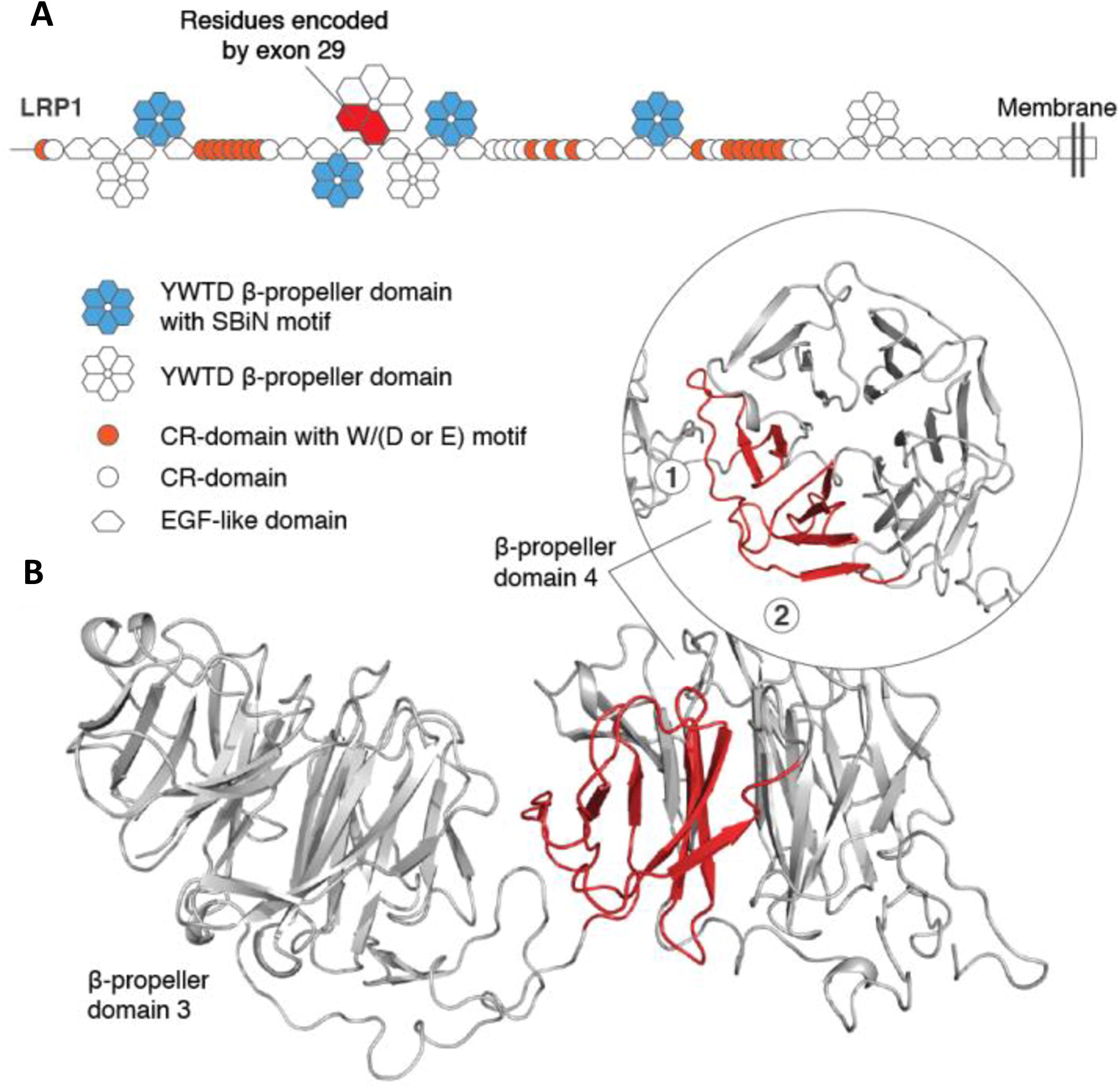
a) Schematic representation of LRP1 domains with the β-propellers as hexagons. The region encoded by exon 29 in the β-propeller 4 is in red. b) Cartoons model for the LRP1 domain β-propeller 3 and 4 obtained from the template of the β-propeller domains 1 and 2 of LRP6 (PDB ID 3s94). The skipping of exon 29 led to the removal of the first two blades out of six of the β-propeller 4.

To investigate whether LRP1 is involved in specific ASD networks of protein-protein interactions, we used ingenuity pathway analysis (IPA) with 75 genes that have previously been implicated in ASD. Interestingly, genes strongly associated to ASD such as *SHANK3, FMR1, SYNGAP1* and *GRIN2B* were found in the same network with *LRP1*, being downstream of LRP1 in the signalling pathway (Supplementary material, Fig. S2).

Given findings suggesting the direct involvement of LRP1 in inflammatory response (32–34), we assessed whether the mutated form of LRP1 may compromise the inflammatory response by measuring IL-6, TNFα and IL-10 markers. Lower mRNA levels were found for all three cytokines in the patient’s lymphoblastoid cell line compared to a control (Figure 3). Comparable results were obtained at the protein level for IL-10 and TNFα (Supplementary material, Fig. S3), whereas IL-6 was not detected from the assay. When we treated the lymphoblastoid cell line with lipopolysaccharide (LPS) to trigger an inflammatory response, a physiological pro-inflammatory response of IL-6 was observed in the control, but not in the patient cell line (Figure 3).

**Figure 3:**
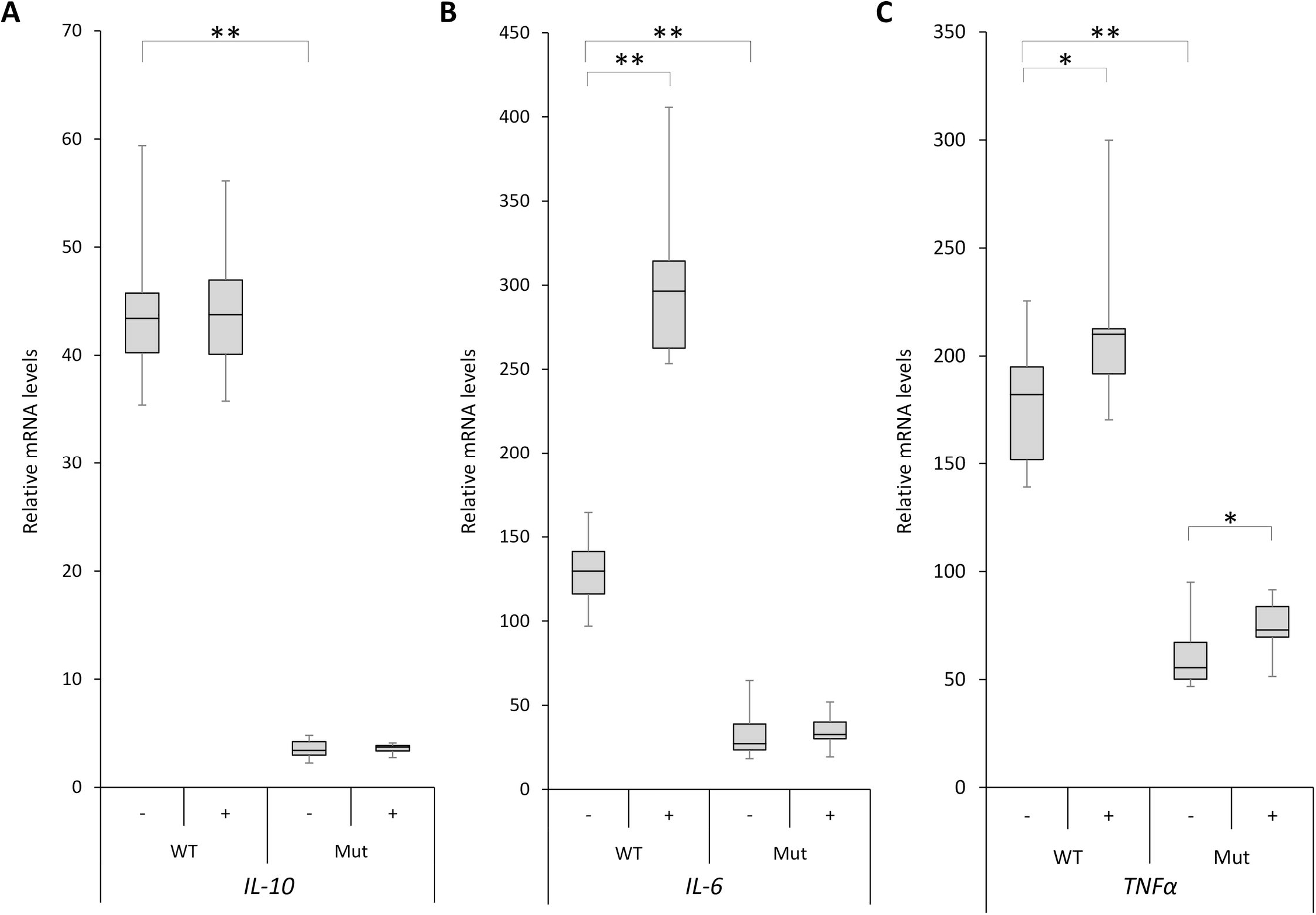
The Boxplots show the cytokines expression in immortalized lymphocyte cell lines from the patient (Mut) and a control (WT) with (+) or without (−) treatment of lipopolysaccharide (LPS) (6h at1μg/mL). mRNA quantification of *IL-10* (A), *IL-6* (B) and *TNF*α (C) were normalized using *ACTB* as endogenous reference. **, *P*<0.0005; *, *P*<0.05.

*LRP1* is ubiquitously expressed, and found in all brain tissues, especially in the cerebellum (Supplementary material, Fig. S4A). Expression reported in BrainCloud and HBT databases shows higher or increasing expression of the gene during foetal or postnatal development, and a relatively stable expression over the rest of the development in all brain tissues (Supplementary material, Fig. S4B-C).

### Involvement of LRP1 in psychiatric disorders by comprehensive analysis of large datasets

Genome-wide significant associations in schizophrenia have recently been reported for SNPs in the *LRP1* region: rs324017 (*P* = 2.12e−08 including replication), rs12814239 (*P* = 1.48e−09), and rs12826178 (*P* = 2.02e−12 including replication) (Supplementary material, Fig. S5) (35). Two of them, rs12814239 (p.C1261C), a synonymous variant in *LRP1*, and rs12826178 (intergenic) are in linkage disequilibrium (D’ = 0.95; r^2^ = 0.74, Caucasians in 1,000 Genomes data) (Supplementary material, Fig. S5). BrainCloud and SMRI Neuropathology Consortium datasets were used to assess potential effects of schizophrenia risk alleles on expression of *LRP1*, but rs12814239 and rs12826178 were not directly genotyped in these datasets, nor other SNPs which would serve as reasonable surrogates (r^2^>0.7).

The identification of a functional DN variant in the *LRP1* gene in an ASD family and the reported associations across the *LRP1* locus in schizophrenia, prompted us to explore the impact of common, rare and DN variants of this gene in several large psychiatric datasets.

The *NPdenovo* and *denovo-db* databases report *de novo* variants in *LRP1* in psychiatric disease projects (supplementary material, Table S5). These DN variants include three highly pathogenic variants, all absent from gnomAD database: a stop mutation from a schizophrenia patient (p.Y2200*), a frame-shift from an ID patient (p.Q3380Sfs*72), and an exon 29 variant in an ASD patient, which is likely to disrupt an exonic splicing enhancer (ESE) site with an effect potentially similar to that reported in patient SJD_33.3. *NPdenovo* data showed association of DN in *LRP1* with ASD (P = 0.039), ID (*P* = 0.008) and SCZ (*P* = 0.008).

We also explored the possible contribution of common variants in *LRP1* by performing a gene-based association study using summary statistics from PGC GWAS data in European populations, which suggests common variants in *LRP1* increase risk of schizophrenia (Gene-based *P* = 6.6E-07), and a trend for ADHD and bipolar disorder, but not in the ASD sample (Table 3). A meta-analysis combining data from six psychiatric disorders strongly implicates *LRP1* common variants across these conditions (*P* = 8.1E-05) (Table 3).

**Table 3.** Results of the gene-based association test of LRP1 across several psychiatric disorders using summary statistics of the PGC data sets.

| **Disease** | **Cases – Controls** | **SNPs tested** | ***P*-value** | **Top SNP** | **Top SNP *P*-value** |
| --- | --- | --- | --- | --- | --- |
| ADHD<sup>a</sup> | 19,099 - 34,194 | 112 | 0.102 | rs34577247 | 0.0023 |
| AN<sup>a</sup> | 3,495 - 10,982 | 149 | 0.406 | rs2228187 | 0.0078 |
| ASD<sup>a</sup> | 6,197 - 7,377 | 80 | 0.99 | rs11172113 | 0.25 |
| BD<sup>a</sup> | 20,352 - 31,358 | 219 | 0.065 | rs10467125 | 0.0014 |
| MDD<sup>b</sup> | 9,240 - 9,519 | 28 | 0.83 | rs1800141 | 0.285 |
| OCD<sup>a</sup> | 2,688 - 7,037 | 137 | 0.36 | rs7398375 | 0.0033 |
| SCZ<sup>a</sup> | 33,640 - 43,456 | 227 | *6.6E-07* | rs12814239 | *2.91E-09* |
| Meta-analysis<sup>a</sup> |  |  | *8.1E-05* |  |  |

Abbreviations: ADHD, attention-deficit/hyperactive disorder; AN, anorexia nervosa; ASD, Autism spectrum disorder; BD, bipolar disorder; MDD, major depressive disorder; OCD, obsessive compulsive disorder; SCZ, schizophrenia;^a^, European individuals from the PGC2 data sets;^b^, PGC1.

A burden analysis using a combined multivariate and collapsing method (CMC) was performed to assess the impact of predicted pathogenic rare or ultra-rare variants (URVs, MAF<0.0001) of *LRP1* in schizophrenia (Swedish case-control) and in the Autism (BCM case-control) data sets. No differences were found in schizophrenia (*P* = 0.63), but a significant burden was observed in autism probands (*P* = 0.048). When data for each phenotype were combined (7,875 control individuals, 6,135 schizophrenic patients and 1,778 ASD probands) no difference in the URVs were observed between schizophrenic patients and controls (*P* = 0.52), whereas a higher burden was observed in ASD patients (*P* = 1.2E-05) (Table 4 and supplementary material, Table S6).

**Table 4.**
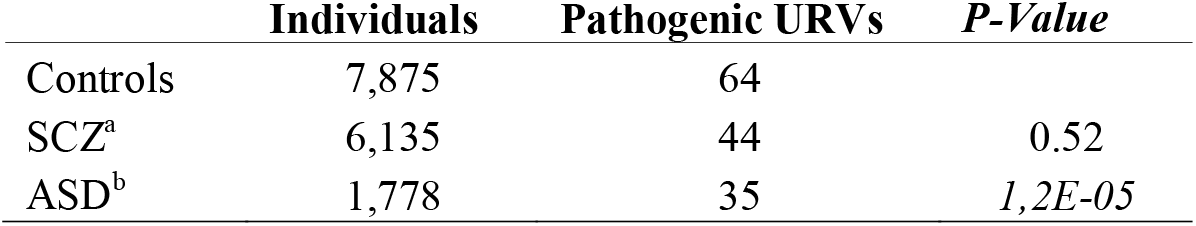
*LRP1* burden analysis of ultra-rare variants in autism spectrum disorder and sc
hizophrenia. Number of total individuals used across different sequencing data sets and number of ultra-rare variants (URVs) predicted to be pathogenic. The selection of variants includes missense, truncating variants and splice site variants (full list of URVs in supplemental Table 9).

Notes:^a^, WES from the Sweden-Schizophrenia population-based Case-Control (dbGAP accession: phs000473.v2.p2);^b^, ARRA Autism Sequencing Collaboration (dbGAP accession: phs000298.v3.p2).

## Discussion

Recent sequencing studies have strongly implicated *de novo* and inherited pathogenic rare variants in ASD (19–29). However, these variants seem to have a varying impact across the heterogeneous ASD phenotype. While the contribution of DN truncating variants has been extensively described in ASD probands with lower IQ, insufficient data are available in regards to inherited truncating variants. A previous study from our group found a higher burden for co-inherited truncating variants in ASD sib-pairs (28), and the burden of these disruptive alleles correlated negatively with non-verbal IQ (NVIQ) (28). Another study showed a significant impact for inherited gene disrupting variants in ASD probands with lower than average IQ (<100) compared with unaffected siblings (29). In the present study we have assessed the impact of rare inherited truncating variants by comparing those that were transmitted vs non-transmitted in singleton families with high function autism (HFA) to establish whether highly disruptive variants have a role in all cases of ASD regardless their comorbidity with intellectual disability. Our results showed no preferential burden transmission from parents to HFA probands, suggesting that truncating alleles may not have a major role across the entire disease spectrum, but may be restricted to those ASD patients with ID.

Correlation between severe disrupting mutations and severe phenotypes has also been reported for other psychiatric diseases: a higher rate of rare truncating variants was found in schizophrenia patients with intellectual disability (36), and in bipolar disorder the burden of inherited truncating alleles was negatively correlated with early age of onset (31). Rare CNV or ultra-rare disruptive alleles were also found associated with lower educational attainment in the general population (37, 38), suggesting that this class of genetic variants may negatively modulate cognitive functions. In summary, all these findings suggest that there is a correlation between the burden of severe mutations and the severity of the disease phenotype.

In our study we also performed enrichment analyses in the list of genes bearing transmitted truncating alleles and DN in HFA probands. No enrichments were observed when we considered this pool of genes, but interestingly, *LRP1* (low-density lipoprotein receptor-related protein-1) recurrently appeared in all psychiatric-related gene-sets. The DN variant found in a HFA patient in *LRP1* is not present in the gnomAD database (138,632 sequenced individuals) and is predicted to disrupt an acceptor splice site. The consequence of this mutation at RNA level is an in-frame skipping of exon 29, which removes two out of six radial blades of the β-propeller 4 of LRP1. It is likely that the β-propeller 4 is not folded in a functional canonical domain, potentially compromising interactions at this and adjacent 3 and 5 β-propellers.

We then investigated the possible involvement of this gene in several psychiatric disorders by exploring a number of genetic datasets. We identified an association of DN variants in *LRP1* with autism and schizophrenia, an association of common variants at gene level in schizophrenia plus an association in a meta-analysis of six psychiatric disorders, and a significant impact for ultra-rare pathogenic variants in ASD. These results implicate common and rare variants in *LRP1* gene across several psychiatric phenotypes. Interestingly, although *LRP1* is ubiquitously expressed, higher expression is found in postnatal cerebral stages, when abnormalities in brain are observed in autistic patients (39). Studies in mice also suggested a peak of *LRP1* expression during early postnatal brain development in several populations of cells including radial glia, immature and mature neurons, microglia and astrocytes (40).

LRP1 has a dual role, being involved in endocytosis and signal transduction, and it binds approximately 40 extracellular ligands mediating a multitude of physiological processes (41). The DN mutation found in a HFA proband in this study affects the structure of a β-propeller domain, which may impair ligand dissociation and the formation of early endosomes (42); the high burden of damaging ultra-rare variants found in autism may abolish the interactions with some of the numerous ligands; the common variants found associated with schizophrenia may potentially exert their role in gene regulation, acting on the antisense RNA *LRP1-AS* transcribed from the *LRP1* locus that negatively regulates *LRP1* expression (43), or regulating alternative transcripts such as the recently identified truncated spliced form of *LRP1* (*smLRP1*) (44).

Considering the high number of ligands and pathways mediated by *LRP1*, it is difficult to pinpoint a specific compromised pathophysiological process. However, several plausible hypotheses can be formulated in relation to its impact on postsynaptic complexes, its role in inflammatory response, insulin signalling, and lipid homeostasis. Firstly, *LRP1* encodes a PSD protein, and plays a role in synaptic integrity and function at the post-synapses by regulating GRIA1 (45), implicated in learning disabilities and autism through *de novo* gain-of-function mutations causing constitutive calcium-channel opening (21). We found *LRP1* in the same network of well-established ASD genes such as *SHANK3, GRIN2B* and *SYNGAP1*. The role of PSD genes in psychiatry has solid evidence (24, 31, 46). Secondly, *LRP1* may exert effects via a compromised inflammatory response. LRP1 regulates inflammation through JNK and NF-kB pathways and has a neuroprotective role in microglia (32–34). After inflammatory insult, the lymphoblastoid cell line of the patient carrying the functional *LRP1* splice variant was not responsive in expressing IL-6 cytokine, suggesting that this pathway may be compromised (47). Thirdly, *LRP1* may mediate insulin signalling in brain, forming a complex with the insulin receptor β (IRβ), and regulating insulin signalling and glucose homeostasis in brain (48). Both processes are involved in synaptic plasticity, memory and learning. Interestingly insulin-related signalling at dendritic spines was implicated in obsessive-compulsive disorder (49) and a stop mutation in the X-linked brain-expressed *IRS4* gene (insulin substrate receptor 4) segregated in schizoaffective patients in an extended family (31). Finally, LRP1 may exert its effects via impairment of lipid homeostasis. Knockout mice showed that neuronal LRP1 is critical for cholesterol and lipid metabolism, and its defect leads to dendritic spine degeneration, synapse loss and neuroinflammation (50). However, it is likely that etiologic genetic variants of *LRP1* exert their role in psychiatric diseases by concurrently impairing more than one pathway at the same time.

It is noteworthy that other lipoprotein receptors from the same family such as *LRP2, LRP1B* and *LRP8* have previously been implicated in autism and psychosis (51–53), suggesting an emerging role for this gene family in psychiatric disorders.

In conclusion, considering our previous and current results, and additional sequencing findings we provide evidence of a relationship between severe mutations and severe ASD phenotype: the accumulation of inherited truncating mutations lead to a more severe phenotype in autism, whereas their impact in high cognitive patients would be limited. Furthermore, we show through a comprehensive analysis of DN, common and rare variants that *LRP1* is a candidate gene with pleiotropic effects across multiple psychiatric phenotypes.

## Materials and Methods

### Selection of ASD subjects and phenotypic assessment

From our collection of ASD families (54, 55), we selected 20 singleton Spanish families without any other psychiatric history amongst relatives. All probands had high functioning autism (HFA), which defined as having full scale IQ greater than 70 (IQ average = 100, SD±14.7, range 80–135). Clinical description is provided in the supplementary material, Table S7.

### Exome sequencing and WES-based genetic relatedness analysis

The sequenced sample included 60 individuals (40 parents and 20 ASD probands). Exome enrichment was performed on 3 μg of genomic DNA extracted from blood, and the exome libraries were applied to an Illumina flowcell for paired-end sequencing on a HiSeq2000 instrument (Illumina) using 76-base reads. Detailed bioinformatic analysis is provided in the Supplement. On average, individuals had 82.1% of the target covered by > 10 reads (Supplementary material, Table S8). Familial relationships were confirmed by genome-wide Identity By Descendent (IBD) analysis in PLINK (56), using WES-derived genotypes (details in the Supplement).

### Variant selection: Rare truncating alleles and de novo variants

Rare variants were defined as those having a minor allele frequency (MAF) <1% in dbSNP135. The truncating alleles selected included nonsense, indels leading to frame-shift, variants in canonical splice sites and start-lost changes. For each family truncating alleles both transmitted to the ASD proband and non-transmitted (i.e. only in parents) were considered. We also examined *de novo* (DN) variants. Rare truncating alleles and DN variants were all validated (N = 263) by Sanger sequencing. During variant validation, we included also 9 unaffected siblings from 8 families to assess the transmission of inherited truncating alleles from parents.

### Enrichment analyses

Enrichment of genes carrying inherited truncating alleles and DN variants were tested against all Gene Ontology (GO) categories and KEGG pathways. Enrichment analysis were also performed using the same pool of genes against several gene sets potentially related to ASD, namely: genes encoding postsynaptic density proteins (PSD) (57), fragile-X mental retardation protein (FMRP) targets (58), *de novo* variants previously found in autism and schizophrenia (30). Both analyses were performed by: *i)* matching genes carrying potentially truncating or DN variants to genes randomly drawn from the genome, after approximate matching on exome enriched coding-sequence length and genic constraint missense Z-score (http://exac.broadinstitute.org) (59); and *ii)* calculating an empirical p-value for observed data for each functional category, using a null distribution of overlap counts from 1,000 randomly drawn gene sets, as described previously (31).

### Effect of LRP1 de novo mutation on splicing

Functional predictions for the *de novo* splice site variant in *LRP1* (chr12:57573110A/G) were performed using three tools: MaxEntScan (60), SPANR (61) and Human Splicing Finder (HSF) (62). To assess functional consequences at RNA level we used cDNA from peripheral blood mononuclear cells (PBMC) of the SJD_33.3 ASD patient (details in the Supplement). PCR products from primers designed in exon 28 and 30 were separated by 10% polyacrylamide gel electrophoresis, and the intensity of the 2 resulting bands measured by a semi-quantitative method (details in the Supplement).

### Protein modelling of LRP1: Consequence of the in-frame skipping of exon 29

The LRP1 wild-type protein was modelled using the Swiss-model platform (63, 64) that was enquired to search homologous templates using the *LRP1* full length sequence (details in the Supplement). The selected model was generated from template 3s94.1.A, corresponding to the structure of the beta-propeller domains 1 and 2 of the human LRP6 (PDB ID 3s94) (65). The LRP1 protein model that includes exon 29 was examined using both SWISS-MODEL and the Robetta server (66, 67).

### LRP1 gene network analysis

We investigated whether *LRP1* was included in a network of genes previously implicated in ASD using Ingenuity Pathway Analysis (IPA) software ((www.ingenuity.com). We computed the most likely network of interactions given a pool of 75 genes with a high probability of involvement in ASD selected from the SFARI database (categories S and 1) (https://gene.sfari.org/database/gene-scoring/).

### Effect of the LRP1 mutated form on inflammatory biomarkers

RNA was extracted from patient SJD_33.3 and a control lymphoblastoid cell line and cDNA was prepared (details in Supplement). We performed quantitative Real-Time PCR (qPCR) and ELISA for the following cytokines: IL-6 and TNFα (pro-inflammatory response), and IL-10 (anti-inflammatory response). Details of these experiments and primers are provided in the Supplementary material, Table S9.

### LRP1: de novo, common and rare variant analyses in psychiatric diseases

Two databases for *de novo* variants were used to identify previous DN in *LRP1* (68, 69). *NPdenovo* was used to assess the overall DN association between *LRP1* and several neuropsychiatric diseases: ASD (6,118 families), SCZ: (1,164 families), epilepsy (647 families) and ID (1,101 families) (68).

Gene-level association for common variants and meta-analysis in *LRP1* was calculated with MAGMA (70) (details in the Supplement), using data sets of European descent only, derived from summary statistics of the Psychiatric Genomics Consortium GWAS https://med.unc.edu/pgc/results-and-downloads) (18, 35, 71–75).

The *LRP1* SNPs significantly associated with schizophrenia in the last GWAS (rs12814239 and rs12826178) (35) were investigated for their effect on *LRP1* expression using the Stanley Medical Research Institute (SMRI) Neuropathology Consortium and BrainCloud data (details in the Supplement). Also, expression of *LRP1* across different developmental stages in human brain was assessed using three data sources: i) Genotype-Tissue Expression project (GTEx) (76); ii) The Human Brain Transcriptome (HBT) (77); and iii) BrainCloud (78).

The analysis of rare variants in *LRP1* was performed using sequencing data of schizophrenia, ASD and control cohorts (details in the Supplement). The selection of potentially etiologic variants is described in the Supplement. A burden analysis was first performed using RVTESTS (79), only in those datasets containing both cases and controls from the same sequencing platform and project (Swedish schizophrenia case-control and BCM autism case-control datasets). A chi-square statistic was then used to compare separately the schizophrenia patient sample (6,135 cases) and combined ASD data-sets (1,778 cases) with the combined control datasets (7,875 individuals).

## Acknowledgments

We are grateful to all patients and their families for their kind participation. We thank all the clinical collaborators that contributed to the diagnosis of the probands. We would also thank the CIBERER Biobank ((http://www.ciberer-biobank.es/), Rafael Valdés-Mas and Xose S. Puente from the University of Oviedo (Spain) for support in bioinformatics analyses, and Barbara Toson (NeuRA) for support in statistical analyses. Exome sequencing services were provided by the National Centre for Genomic Analysis (CNAG).

**Abbreviations**
ADHD: attention-deficit hyperactivity disorder
ASD: Autism spectrum disorder
BD: bipolar disorder
CMC: combined multivariate and collapsing method
DN: *de novo* variant
FMRP: Fragile X mental retardation protein target genes
GTEx: Genotype-Tissue Expression project
HBT: The Human Brain Transcriptome
HFA: high functioning autism
ID: intellectual disability
IPA: ingenuity pathway analysis
IQ: intelligence quotient
MAF: minor allele frequency
MDD: major depressive disorder
NVIQ: non-verbal IQ
PSD: Post-Synaptic Density
SCZ: schizophrenia
SNV: single nucleotide variation
URV: ultra-rare variants
WES: whole-exome sequencing
WGS: whole-genome sequencing studies

## References

1 Elsabbagh, M., Divan, G., Koh, Y.J., Kim, Y.S., Kauchali, S., Marcin, C., Montiel-Nava, C., Patel, V., Paula, C.S., Wang, C. et al. (2012) Global prevalence of autism and other pervasive developmental disorders. Autism research: official journal of the International Society for Autism Research, 5, 160–179.

2 Freitag, C.M. (2007) The genetics of autistic disorders and its clinical relevance: a review of the literature. Molecular psychiatry, 12, 2–22.

3 Idring, S., Lundberg, M., Sturm, H., Dalman, C., Gumpert, C., Rai, D., Lee, B.K. and Magnusson, C. (2015) Changes in prevalence of autism spectrum disorders in 2001–2011: findings from the Stockholm youth cohort. Journal of autism and developmental disorders, 45, 1766–1773.

4 Vivanti, G., Barbaro, J., Hudry, K., Dissanayake, C. and Prior, M. (2013) Intellectual development in autism spectrum disorders: new insights from longitudinal studies. Frontiers in human neuroscience, 7, 354.

5 Sandin, S., Lichtenstein, P., Kuja-Halkola, R., Hultman, C., Larsson, H. and Reichenberg, A. (2017) The Heritability of Autism Spectrum Disorder. Jama, 318, 1182–1184.

6 Betancur, C. (2011) Etiological heterogeneity in autism spectrum disorders: more than 100 genetic and genomic disorders and still counting. Brain research, 1380, 42–77.

7 Ronemus, M., Iossifov, I., Levy, D. and Wigler, M. (2014) The role of de novo mutations in the genetics of autism spectrum disorders. Nature reviews. Genetics, 15, 133–141.

8 Young, A.M., Chakrabarti, B., Roberts, D., Lai, M.C., Suckling, J. and Baron-Cohen, S. (2016) From molecules to neural morphology: understanding neuroinflammation in autism spectrum condition. Molecular autism, 7, 9.

9 Marazziti, D., Baroni, S., Picchetti, M., Landi, P., Silvestri, S., Vatteroni, E. and Catena Dell’Osso, M. (2012) Psychiatric disorders and mitochondrial dysfunctions. European review for medical and pharmacological sciences, 16, 270–275.

10 Anitha, A., Nakamura, K., Thanseem, I., Yamada, K., Iwayama, Y., Toyota, T., Matsuzaki, H., Miyachi, T., Yamada, S., Tsujii, M. et al. (2012) Brain region-specific altered expression and association of mitochondria-related genes in autism. Molecular autism, 3, 12.

12 Wang, X., Kery, R. and Xiong, Q. (2017) Synaptopathology in autism spectrum disorders: Complex effects of synaptic genes on neural circuits. Progress in neuro-psychopharmacology & biological psychiatry, in press.

12 Anney, R., Klei, L., Pinto, D., Regan, R., Conroy, J., Magalhaes, T.R., Correia, C., Abrahams, B.S., Sykes, N., Pagnamenta, A.T. et al. (2010) A genome-wide scan for common alleles affecting risk for autism. Human molecular genetics, 19, 4072–4082.

13 Weiss, L.A., Arking, D.E., Gene Discovery Project of Johns, H., the Autism, C., Daly, M.J. and Chakravarti, A. (2009) A genome-wide linkage and association scan reveals novel loci for autism. Nature, 461, 802–808.

14 Wang, K., Zhang, H., Ma, D., Bucan, M., Glessner, J.T., Abrahams, B.S., Salyakina, D., Imielinski, M., Bradfield, J.P., Sleiman, P.M. et al. (2009) Common genetic variants on 5p14.1 associate with autism spectrum disorders. Nature, 459, 528–533.

15 Torrico, B., Chiocchetti, A.G., Bacchelli, E., Trabetti, E., Hervas, A., Franke, B., Buitelaar, J.K., Rommelse, N., Yousaf, A., Duketis, E. et al. (2017) Lack of replication of previous autism spectrum disorder GWAS hits in European populations. Autism research: official journal of the International Society for Autism Research, 10, 202–211.

16 Gaugler, T., Klei, L., Sanders, S.J., Bodea, C.A., Goldberg, A.P., Lee, A.B., Mahajan, M., Manaa, D., Pawitan, Y., Reichert, J. et al. (2014) Most genetic risk for autism resides with common variation. Nature genetics, 46, 881–885.

17 Cross-Disorder Group of the Psychiatric Genomics, C., Lee, S.H., Ripke, S., Neale, B.M., Faraone, S.V., Purcell, S.M., Perlis, R.H., Mowry, B.J., Thapar, A., Goddard, M.E. et al. (2013) Genetic relationship between five psychiatric disorders estimated from genome-wide SNPs. Nature genetics, 45, 984–994.

18 Autism Spectrum Disorders Working Group of The Psychiatric Genomics, C. (2017) Meta-analysis of GWAS of over 16,000 individuals with autism spectrum disorder highlights a novel locus at 10q24.32 and a significant overlap with schizophrenia. Molecular autism, 8, 21.

19 Yuen, R.K., Thiruvahindrapuram, B., Merico, D., Walker, S., Tammimies, K., Hoang, N., Chrysler, C., Nalpathamkalam, T., Pellecchia, G., Liu, Y. et al. (2015) Whole-genome sequencing of quartet families with autism spectrum disorder. Nature medicine, 21, 185–191.

20 RK, C.Y., Merico, D., Bookman, M., J, L.H., Thiruvahindrapuram, B., Patel, R.V., Whitney, J., Deflaux, N., Bingham, J., Wang, Z. et al. (2017) Whole genome sequencing resource identifies 18 new candidate genes for autism spectrum disorder. Nature neuroscience, 20, 602–611.

21 Geisheker, M.R., Heymann, G., Wang, T., Coe, B.P., Turner, T.N., Stessman, H.A.F., Hoekzema, K., Kvarnung, M., Shaw, M., Friend, K. et al. (2017) Hotspots of missense mutation identify neurodevelopmental disorder genes and functional domains. Nature neuroscience, 20, 1043–1051.

22 O’Roak, B.J., Vives, L., Girirajan, S., Karakoc, E., Krumm, N., Coe, B.P., Levy, R., Ko, A., Lee, C., Smith, J.D. et al. (2012) Sporadic autism exomes reveal a highly interconnected protein network of de novo mutations. Nature, 485, 246–250.

23 Iossifov, I., O’Roak, B.J., Sanders, S.J., Ronemus, M., Krumm, N., Levy, D., Stessman, H.A., Witherspoon, K.T., Vives, L., Patterson, K.E. et al. (2014) The contribution of de novo coding mutations to autism spectrum disorder. Nature, 515, 216–221.

24 De Rubeis, S., He, X., Goldberg, A.P., Poultney, C.S., Samocha, K., Cicek, A.E., Kou, Y., Liu, L., Fromer, M., Walker, S. et al. (2014) Synaptic, transcriptional and chromatin genes disrupted in autism. Nature, 515, 209–215.

25 Kosmicki, J.A., Samocha, K.E., Howrigan, D.P., Sanders, S.J., Slowikowski, K., Lek, M., Karczewski, K.J., Cutler, D.J., Devlin, B., Roeder, K. et al. (2017) Refining the role of de novo protein-truncating variants in neurodevelopmental disorders by using population reference samples. Nature genetics, 49, 504–510.

26 Neale, B.M., Kou, Y., Liu, L., Ma’ayan, A., Samocha, K.E., Sabo, A., Lin, C.F., Stevens, C., Wang, L.S., Makarov, V. et al. (2012) Patterns and rates of exonic de novo mutations in autism spectrum disorders. Nature, 485, 242–245.

27 Sanders, S.J., He, X., Willsey, A.J., Ercan-Sencicek, A.G., Samocha, K.E., Cicek, A.E., Murtha, M.T., Bal, V.H., Bishop, S.L., Dong, S. et al. (2015) Insights into Autism Spectrum Disorder Genomic Architecture and Biology from 71 Risk Loci. Neuron, 87, 1215–1233.

28 Toma, C., Torrico, B., Hervas, A., Valdes-Mas, R., Tristan-Noguero, A., Padillo, V., Maristany, M., Salgado, M., Arenas, C., Puente, X.S. et al. (2014) Exome sequencing in multiplex autism families suggests a major role for heterozygous truncating mutations. Molecular psychiatry, 19, 784–790.

29 Krumm, N., Turner, T.N., Baker, C., Vives, L., Mohajeri, K., Witherspoon, K., Raja, A., Coe, B.P., Stessman, H.A., He, Z.X. et al. (2015) Excess of rare, inherited truncating mutations in autism. Nature genetics, 47, 582–588.

30 Fromer, M., Pocklington, A.J., Kavanagh, D.H., Williams, H.J., Dwyer, S., Gormley, P., Georgieva, L., Rees, E., Palta, P., Ruderfer, D.M. et al. (2014) De novo mutations in schizophrenia implicate synaptic networks. Nature, 506, 179–184.

31 Toma, C., Shaw, D., Allcock, R., Heath, A., Pierce, K., Mitchell, P., Schofield, P. and Fullerton, J. (2018) An Examination of Multiple Classes of Rare Variants in Extended Families with Bipolar Disorder. Translational Psychiatry, in press.

32 Yang, L., Liu, C.C., Zheng, H., Kanekiyo, T., Atagi, Y., Jia, L., Wang, D., N’Songo, A., Can, D., Xu, H. et al. (2016) LRP1 modulates the microglial immune response via regulation of JNK and NF-kappaB signaling pathways. Journal of neuroinflammation, 13, 304.

33 Mantuano, E., Brifault, C., Lam, M.S., Azmoon, P., Gilder, A.S. and Gonias, S.L. (2016) LDL receptor-related protein-1 regulates NFkappaB and microRNA-155 in macrophages to control the inflammatory response. Proceedings of the National Academy of Sciences of the United States of America, 113, 1369–1374.

34 Chuang, T.Y., Guo, Y., Seki, S.M., Rosen, A.M., Johanson, D.M., Mandell, J.W., Lucchinetti, C.F. and Gaultier, A. (2016) LRP1 expression in microglia is protective during CNS autoimmunity. Acta neuropathologica communications, 4, 68.

35 Schizophrenia Working Group of the Psychiatric Genomics, C. (2014) Biological insights from 108 schizophrenia-associated genetic loci. Nature, 511, 421–427.

36 Purcell, S.M., Moran, J.L., Fromer, M., Ruderfer, D., Solovieff, N., Roussos, P., O’Dushlaine, C., Chambert, K., Bergen, S.E., Kahler, A. et al. (2014) A polygenic burden of rare disruptive mutations in schizophrenia. Nature, 506, 185–190.

37 Ganna, A., Genovese, G., Howrigan, D.P., Byrnes, A., Kurki, M., Zekavat, S.M., Whelan, C.W., Kals, M., Nivard, M.G., Bloemendal, A. et al. (2016) Ultra-rare disruptive and damaging mutations influence educational attainment in the general population. Nature neuroscience, 19, 1563–1565.

38 Mannik, K., Magi, R., Mace, A., Cole, B., Guyatt, A.L., Shihab, H.A., Maillard, A.M., Alavere, H., Kolk, A., Reigo, A. et al. (2015) Copy number variations and cognitive phenotypes in unselected populations. Jama, 313, 2044–2054.

39 Donovan, A.P. and Basson, M.A. (2017) The neuroanatomy of autism – a developmental perspective. Journal of anatomy, 230, 4–15.

40 Auderset, L., Cullen, C.L. and Young, K.M. (2016) Low Density Lipoprotein-Receptor Related Protein 1 Is Differentially Expressed by Neuronal and Glial Populations in the Developing and Mature Mouse Central Nervous System. PloS one, 11, e0155878.

41 Zlokovic, B.V., Deane, R., Sagare, A.P., Bell, R.D. and Winkler, E.A. (2010) Low-density lipoprotein receptor-related protein-1: a serial clearance homeostatic mechanism controlling Alzheimer’s amyloid beta-peptide elimination from the brain. Journal of neurochemistry, 115, 1077–1089.

42 Jeon, H., Meng, W., Takagi, J., Eck, M.J., Springer, T.A. and Blacklow, S.C. (2001) Implications for familial hypercholesterolemia from the structure of the LDL receptor YWTD-EGF domain pair. Nature structural biology, 8, 499–504.

43 Yamanaka, Y., Faghihi, M.A., Magistri, M., Alvarez-Garcia, O., Lotz, M. and Wahlestedt, C. (2015) Antisense RNA controls LRP1 Sense transcript expression through interaction with a chromatin-associated protein, HMGB2. Cell reports, 11, 967–976.

44 Kolb, M., Kurz, S., Schafer, A., Huse, K., Dietz, A., Wichmann, G. and Birkenmeier, G. (2017) Verification and characterization of an alternative low density lipoprotein receptor-related protein 1 splice variant. PloS one, 12, e0180354.

45 Gan, M., Jiang, P., McLean, P., Kanekiyo, T. and Bu, G. (2014) Low-density lipoprotein receptor-related protein 1 (LRP1) regulates the stability and function of GluA1 alpha-amino-3-hydroxy-5-methyl-4-isoxazole propionic acid (AMPA) receptor in neurons. PloS one, 9, e113237.

46 Network and Pathway Analysis Subgroup of Psychiatric Genomics, C. (2015) Psychiatric genome-wide association study analyses implicate neuronal, immune and histone pathways. Nature neuroscience, 18, 199–209.

47 Masi, A., Glozier, N., Dale, R. and Guastella, A.J. (2017) The Immune System, Cytokines, and Biomarkers in Autism Spectrum Disorder. Neuroscience bulletin, 33, 194–204.

48 Liu, C.C., Hu, J., Tsai, C.W., Yue, M., Melrose, H.L., Kanekiyo, T. and Bu, G. (2015) Neuronal LRP1 regulates glucose metabolism and insulin signaling in the brain. The Journal of neuroscience: the official journal of the Society for Neuroscience, 35, 5851–5859.

49 van de Vondervoort, I., Poelmans, G., Aschrafi, A., Pauls, D.L., Buitelaar, J.K., Glennon, J.C. and Franke, B. (2016) An integrated molecular landscape implicates the regulation of dendritic spine formation through insulin-related signalling in obsessive-compulsive disorder. Journal of psychiatry & neuroscience: JPN, 41, 280–285.

50 Liu, Q., Trotter, J., Zhang, J., Peters, M.M., Cheng, H., Bao, J., Han, X., Weeber, E.J. and Bu, G. (2010) Neuronal LRP1 knockout in adult mice leads to impaired brain lipid metabolism and progressive, age-dependent synapse loss and neurodegeneration. The Journal of neuroscience: the official journal of the Society for Neuroscience, 30, 17068–17078.

51 Ionita-Laza, I., Makarov, V., Consortium, A.A.S. and Buxbaum, J.D. (2012) Scan-statistic approach identifies clusters of rare disease variants in LRP2, a gene linked and associated with autism spectrum disorders, in three datasets. American journal of human genetics, 90, 1002–1013.

52 Li, M., Huang, L., Grigoroiu-Serbanescu, M., Bergen, S.E., Landen, M., Hultman, C.M., Forstner, A.J., Strohmaier, J., Hecker, J., Schulze, T.G. et al. (2016) Convergent Lines of Evidence Support LRP8 as a Susceptibility Gene for Psychosis. Molecular neurobiology, 53, 6608–6619.

53 Alliey-Rodriguez, N., Grey, T.A., Shafee, R., Padmanabhan, J., Tandon, N., Klinger, M., Spring, J., Coppes, L., Reis, K., Keshavan, M.S. et al. (2017) Common variants of NRXN1, LRP1B and RORA are associated with increased ventricular volumes in psychosis – GWAS findings from the B-SNIP deep phenotyping study. bioRxiv, in press.

54 Toma, C., Torrico, B., Hervas, A., Salgado, M., Rueda, I., Valdes-Mas, R., Buitelaar, J.K., Rommelse, N., Franke, B., Freitag, C. et al. (2015) Common and rare variants of microRNA genes in autism spectrum disorders. The world journal of biological psychiatry: the official journal of the World Federation of Societies of Biological Psychiatry, in press., 1–11.

55 Torrico, B., Fernandez-Castillo, N., Hervas, A., Mila, M., Salgado, M., Rueda, I., Buitelaar, J.K., Rommelse, N., Oerlemans, A.M., Bralten, J. et al. (2015) Contribution of common and rare variants of the PTCHD1 gene to autism spectrum disorders and intellectual disability. European journal of human genetics: EJHG, 23, 1694–1701.

56 Purcell, S., Neale, B., Todd-Brown, K., Thomas, L., Ferreira, M.A., Bender, D., Maller, J., Sklar, P., de Bakker, P.I., Daly, M.J. et al. (2007) PLINK: a tool set for whole-genome association and population-based linkage analyses. American journal of human genetics, 81, 559–575.

57 Bayes, A., van de Lagemaat, L.N., Collins, M.O., Croning, M.D., Whittle, I.R., Choudhary, J.S. and Grant, S.G. (2011) Characterization of the proteome, diseases and evolution of the human postsynaptic density. Nature neuroscience, 14, 19–21.

58 Darnell, J.C., Van Driesche, S.J., Zhang, C., Hung, K.Y., Mele, A., Fraser, C.E., Stone, E.F., Chen, C., Fak, J.J., Chi, S.W. et al. (2011) FMRP stalls ribosomal translocation on mRNAs linked to synaptic function and autism. Cell, 146, 247–261.

59 Lek, M., Karczewski, K.J., Minikel, E.V., Samocha, K.E., Banks, E., Fennell, T., O’Donnell-Luria, A.H., Ware, J.S., Hill, A.J., Cummings, B.B. et al. (2016) Analysis of protein-coding genetic variation in 60,706 humans. Nature, 536, 285–291.

60 Yeo, G. and Burge, C.B. (2004) Maximum entropy modeling of short sequence motifs with applications to RNA splicing signals. Journal of computational biology: a journal of computational molecular cell biology, 11, 377–394.

61 Xiong, H.Y., Alipanahi, B., Lee, L.J., Bretschneider, H., Merico, D., Yuen, R.K., Hua, Y., Gueroussov, S., Najafabadi, H.S., Hughes, T.R. et al. (2015) RNA splicing. The human splicing code reveals new insights into the genetic determinants of disease. Science, 347, 1254806.

62 Desmet, F.O., Hamroun, D., Lalande, M., Collod-Beroud, G., Claustres, M. and Beroud, C. (2009) Human Splicing Finder: an online bioinformatics tool to predict splicing signals. Nucleic acids research, 37, e67.

63 Arnold, K., Bordoli, L., Kopp, J. and Schwede, T. (2006) The SWISS-MODEL workspace: a web-based environment for protein structure homology modelling. Bioinformatics, 22, 195–201.

64 Biasini, M., Bienert, S., Waterhouse, A., Arnold, K., Studer, G., Schmidt, T., Kiefer, F., Gallo Cassarino, T., Bertoni, M., Bordoli, L. et al. (2014) SWISS-MODEL: modelling protein tertiary and quaternary structure using evolutionary information. Nucleic acids research, 42, W252–258.

65 Cheng, Z., Biechele, T., Wei, Z., Morrone, S., Moon, R.T., Wang, L. and Xu, W. (2011) Crystal structures of the extracellular domain of LRP6 and its complex with DKK1. Nature structural & molecular biology, 18, 1204–1210.

66 Raman, S., Vernon, R., Thompson, J., Tyka, M., Sadreyev, R., Pei, J., Kim, D., Kellogg, E., DiMaio, F., Lange, O. et al. (2009) Structure prediction for CASP8 with all-atom refinement using Rosetta. Proteins, 77 Suppl 9, 89–99.

67 Song, Y., DiMaio, F., Wang, R.Y., Kim, D., Miles, C., Brunette, T., Thompson, J. and Baker, D. (2013) High-resolution comparative modeling with RosettaCM. Structure, 21, 1735–1742.

68 Li, J., Cai, T., Jiang, Y., Chen, H., He, X., Chen, C., Li, X., Shao, Q., Ran, X., Li, Z. et al. (2016) Genes with de novo mutations are shared by four neuropsychiatric disorders discovered from NPdenovo database. Molecular psychiatry, 21, 298.

69 Turner, T.N., Yi, Q., Krumm, N., Huddleston, J., Hoekzema, K., HA, F.S., Doebley, A.L., Bernier, R.A., Nickerson, D.A. and Eichler, E.E. (2017) denovo-db: a compendium of human de novo variants. Nucleic acids research, 45, D804–D811.

70 de Leeuw, C.A., Mooij, J.M., Heskes, T. and Posthuma, D. (2015) MAGMA: generalized gene-set analysis of GWAS data. PLoS computational biology, 11, e1004219.

71 Demontis, D., Walters, R.K., Martin, J., Mattheisen, M., Als, T.D., Agerbo, E., Belliveau, R., Bybjerg-Grauholm, J., Bækved-Hansen, M., Cerrato, F. et al. (2017) Discovery Of The First Genome-Wide Significant Risk Loci For ADHD. bioRxiv, in press.

72 Major Depressive Disorder Working Group of the Psychiatric, G.C., Ripke, S., Wray, N.R., Lewis, C.M., Hamilton, S.P., Weissman, M.M., Breen, G., Byrne, E.M., Blackwood, D.H., Boomsma, D.I. et al. (2013) A mega-analysis of genome-wide association studies for major depressive disorder. Molecular psychiatry, 18, 497–511.

73 Stahl, E., Forstner, A., McQuillin, A., Ripke, S., Ophoff, R., Scott, L., Cichon, S., Andreassen, O.A., Sklar, P., Kelsoe, J. et al. (2017) Genomewide association study identifies 30 loci associated with bipolar disorder. bioRxiv, in press.

74 Duncan, L., Yilmaz, Z., Gaspar, H., Walters, R., Goldstein, J., Anttila, V., Bulik-Sullivan, B., Ripke, S., Eating Disorders Working Group of the Psychiatric Genomics, C., Thornton, L. et al. (2017) Significant Locus and Metabolic Genetic Correlations Revealed in Genome-Wide Association Study of Anorexia Nervosa. The American journal of psychiatry, 174, 850–858.

75 International Obsessive Compulsive Disorder Foundation Genetics, C. and Studies, O.C.D.C.G.A. (2017) Revealing the complex genetic architecture of obsessive-compulsive disorder using meta-analysis. Molecular psychiatry, in press.

76 Consortium, G.T. (2013) The Genotype-Tissue Expression (GTEx) project. Nature genetics, 45, 580–585.

77 Kang, H.J., Kawasawa, Y.I., Cheng, F., Zhu, Y., Xu, X., Li, M., Sousa, A.M., Pletikos, M., Meyer, K.A., Sedmak, G. et al. (2011) Spatio-temporal transcriptome of the human brain. Nature, 478, 483–489.

78 Colantuoni, C., Lipska, B.K., Ye, T., Hyde, T.M., Tao, R., Leek, J.T., Colantuoni, E.A., Elkahloun, A.G., Herman, M.M., Weinberger, D.R. et al. (2011) Temporal dynamics and genetic control of transcription in the human prefrontal cortex. Nature, 478, 519–523.

79 Zhan, X., Hu, Y., Li, B., Abecasis, G.R. and Liu, D.J. (2016) RVTESTS: an efficient and comprehensive tool for rare variant association analysis using sequence data. Bioinformatics, 32, 1423–1426.

